# Soil viral response to drought disturbance and ecosystem recovery in an artificial rainforest

**DOI:** 10.64898/2026.09.28.755132

**Authors:** María Touceda-Suárez, Alise J. Ponsero, Linnea K. Hernandez, Juliana Gil-Loaiza, Jane Fudyma, Meara Clark, Christiane Werner, S. Nemiah Ladd, Malak M. Tfaily, Albert Barberán, Laura K. Meredith

**Author notes:** Corresponding author: Laura K. Meredith.

## Abstract

Anthropogenic climate change is driving increased temperature and altered precipitation patterns in tropical rainforests, yet the impact of severe drought on soil viral communities and their recovery capacity remains poorly understood. Soil viruses can exert strong controls on bacterial populations impacting microbially-mediated soil processes such as carbon cycling. At the same time, viruses are very sensitive to changes in the soil environment. We leverage meta-omics data from soils collected during a two-month drought and one-month recovery experiment carried in the Biosphere 2 Tropical Rainforest artificial biome. Our results indicate that soil viral community structure responds to small scale soil habitat heterogeneity more strongly than on overall changes in moisture. Additionally, the abundance of soil viruses was associated with a higher diversity of metabolites in the soil, suggesting viruses play a key role in soil biogeochemical processes. Altogether, our study suggests that soil viruses are an active part of tropical rainforest soil microbial communities and thus that is necessary to understand their response to drought.

## 1. Introduction

Soil viruses have the potential to control microbial populations through predation (infection and lysis) and metabolic reprogramming, thus changing community structure and microbially mediated soil processes (Thompson et al., 2011; Emerson et al., 2018; Osburn et al., 2024). Particularly, viral lysis can cause the release of organic carbon and nutrients from microbial cells, possibly altering soil biogeochemical cycles (Albright et al., 2022). At the same time, soil viruses are highly sensitive to changes in the soil environment, potentially more so than their bacterial hosts (Cao et al., 2022; Jansson, 2023). Moisture plays an important role in soil viral survival and function, in fact, ecosystem rewetting dynamics affect the structure of the viral community (Coclet et al., 2023), in some cases even when the bacterial community is unchanged (Santos-Medellín et al., 2023). Furthermore, increases in soil moisture correlate with the shift from lysogenic to lytic infection (Wu et al., 2021), and when environmental conditions become unsustainable, in extreme drought for instance, soil viruses could induce host dormancy through the expression of sigma factors (Gabiatti et al., 2018).

Anthropogenic climate change is increasing temperature and changing precipitation patterns (IPCC, 2019). Tropical rainforests have been experiencing, and are projected to experience, increased droughts that threaten to cause critical ecosystem transitions such as forest decline and savannization (Boisier et al., 2015; Carvalho et al., 2020). Due to the importance of rainforests for the global climatic equilibrium (Watson et al., 2018) great efforts have been made to estimate the impacts of drought on their soil biogeochemistry (Davidson et al. 2008; Doughty et al. 2015), greenhouse and trace gas fluxes (Bréchet et al., 2019; Honeker et al., 2023; Pugliese et al., 2023), nutrient pools (O’Connell et al., 2018), and carbon balance (Zhang et al., 2015; Werner et al., 2021). Tropical rainforests store around 25% of global soil carbon stocks, but also have short carbon residence times and highly sensitive carbon balance amplified by high microbial activity (McFarlane et al., 2024). Therefore, unveiling the role of viruses in tropical rainforest soil processes and their response to drought becomes increasingly important.

Current theory suggests that high functional diversity in microbial communities and diverse microbial trophic networks could support higher ecosystem resilience (Shade et al., 2012). However, biotic interactions can produce cascading or even counterintuitive effects that make system resilience hard to predict (Philippot et al., 2021). For example, population dynamics of soil microorganisms responded differently to drought depending on the organism and ecosystem in question, and the strength and duration of the disturbance (Bardgett and Caruso 2020). This underscores the need for ecosystem-specific research to uncover the contribution of underexplored fractions of the microbial communities in ecosystem resilience. Given their role in the trophic chain, their influence on community function, and their potential sensibility to changes in the environment, it is important to improve our understanding of the responses and recovery of soil viruses from disturbances such as drought.

In this project, we explored the viral response to drought using meta-omics (metagenomics, metatranscriptomics, and metabolomics) data from soil subjected to a controlled two-month drought and one-month recovery in a large-scale artificial tropical rainforest ecosystem (Werner et al., 2021). We inferred soil viral genomes from metagenomes and metatranscriptomes to analyze the effect of soil moisture changes on soil DNA and RNA viral community structure and activity. Additionally, we examined relationships between viral abundances and metabolite diversity, the oxidative state of carbon (NOSC), and the relative abundance of less metabolically available carbon forms such as lipids to evaluate their role in carbon and nutrient cycling. We hypothesized that: (i) viral community structure and activity would respond to changes in moisture with a decrease in viral diversity, abundance and activity during drought, followed by a partial return to pre-drought levels with the recovery of moisture levels as has been reported for other microorganisms in tropical rainforests (L. Li et al., 2021) and for viruses in other ecosystems (Santos-Medellín et al., 2023); and (ii) there would be an increased differentiation of viral community composition during the drought, followed by a shift towards a community composition similar to pre-drought as moisture levels recovered. Viral lifestyles are potentially dependent on host abundance and activity, for example, viruses with temperate lifestyles could be better suited to cope with long periods of low host activity such as drought (Stewart and Levin 1984). We therefore expected to find (iii) higher abundance and activity of virulent viruses during the high moisture periods (pre-drought and recovery), causing potential increases in lysis that could lead to (iv) the release of nutrients and more metabolically available forms of carbon to the soil. As the first study to investigate changes in soil viral communities and their association with soil processes in a tropical ecosystem under severe drought, our results help reveal the virus-mediated impacts of these increasingly prevalent conditions on the functioning of tropical ecosystems.

## 2. Materials and methods

### 2.1. Experimental design, sample collection and soil water measurement

The drought experiment was carried out in the Biosphere 2 Tropical Rainforest. Biosphere 2 is a 12,700 m^2^ steel and glass-enclosed building located in Oracle, Arizona, USA with the ability to control rainfall inside. This 30-year-old artificial tropical rainforest harbors approximately 70 species of trees and shrubs that form a canopy and understory distributed across a range of variable topography and microhabitats to represent biogeochemical cycles present in natural rainforests (Leigh et al., 1999). In late 2019, a drought experiment was conducted as part of the Water, Atmosphere and Life Dynamics (WALD) campaign (Werner et al., 2021). Briefly, the regular rain precipitation scheme of three rain events per week was interrupted on October 6^th^, 2019, to subject the forest to a 67-day long drought during which pre-drought temperature was maintained between 20 and 26.7 °C. The first rainfall simulated event after drought took place on December 12^th^, 2019, by spraying 15,000 L of irrigation water from the top of the Biosphere 2 tropical rainforest. A second rainfall event was simulated seven days later, on December 19^th^, 2019, followed by a reintroduction of the regular rain precipitation scheme. A list of the drought events simulated in this system can be found in Supplementary Table 1. Soil samples were collected in quadruplicate at two adjacent sites (Sites 1 and 2 as depicted in Supplementary Figure 1C, Fig. S2 in Werner et al., 2019, and Fig. S1 in Honeker et al., 2023) in the lowland region of the Biosphere 2 tropical rainforest. Sampling locations were randomly pre-assigned from within a set of four 1-m x 1-m quadrants each site, leaving at least 20-cm between each sample and 20-cm distance from trees. Plant species composition of each site can be found in Supplementary Table 2; a description of the soil can be found in Supplementary Methods. Soil was collected using a large auger (1-m long by 3.04 cm diameter) before the interruption of rain (Pre-drought, PD: October 9^th^ 2019) and in middle stages of drought (Mid-drought, MD: November 29^th^ 2019), and using a hand auger (30 cm in length and 2.2 cm diameter) immediately prior to the first rewet event (Late-drought, LD; December 12^th^ 2019), at 3 hours, 24 hours, and 48 hours after the rewet event (RW1: December 12^th^ 2019; RW2: December 13^th^ 2019; and RW3: December 14^th^ 2019), and at the start and end of the recovery phase (R1: December 19^th^ 2019; R2: January 2^nd^ 2020). Core samples were stored on ice and immediately brought to the lab where soil from the 0-15 cm depths was separated from other depths, most roots were removed by hand, and 1 g of bulk soil was stored in Lifeguard® Soil Preservation Solution (Qiagen N.V., Venlo, Netherlands) and stored at -80°C for RNA/DNA extraction. See Supplementary Figure 1A for a schematic representation of the experiment phases and sample collection timepoints. Soil moisture content measurements were obtained using a portable probe and LabQuest viewer (Vernier). Soil water matric potential were collected from environmental sensors as described in Werner et al. 2021 (see also for more detailed description of B2WALD campaign methodologies).

### 2.2. Fourier transform ion-coupled resonance mass spectrometry (FTICR-MS)

We performed water extractions followed by solid phase extraction (SPE) to characterize soil metabolites, as water is the primary solvent in natural soil systems and best represents environmentally relevant conditions. This approach enables extraction of polar and semi-polar compounds that are most likely to be bioavailable to soil microorganisms. Water extraction procedures followed established protocols for bulk metabolite characterization (Tfaily et al., 2015). Extracted samples were analyzed at Pacific Northwest National Laboratory (PNNL) using a 12-Tesla Bruker fourier transform ion-coupled resonance mass spectrometry (FTICR-MS) for high-resolution characterization of soil organic matter metabolites as detailed in Honeker et al. 2023). Putative chemical formulae were assigned using Formularity (Tolić et al., 2017) with the following criteria: S/N > 7, mass measurement error <1 ppm and taking into consideration the presence of C, H, O, N, S and P and excluding other elements (Tfaily et al., 2018). To ensure consistent formula assignment and eliminate mass shifts, all sample peak lists were aligned to each other, and we ensured that only formulae with the lowest error between predicted and observed m/z and the lowest number of heteroatoms and at least four oxygen atoms for each phosphorus atom were kept (Tfaily et al., 2018). We evaluated the chemical character of peaks in electrospray ionization (ESI) FTICR-MS spectra and calculated lipids relative abundance and the oxidative state of carbon (NOSC) using MetaboDirect (Ayala-Ortiz et al., 2023).

### 2.3. Molecular assays and DNA/RNA extractions and sequencing

DNA and RNA were co-extracted from 1 g of soil using the RNeasy Powersoil Total RNA kit (Qiagen, 12866–25) coupled with the RNeasy Powersoil DNA Elution kit (Qiagen, 12867–25) following the manufacturer’s protocol. Nucleic acid concentrations and quality were measured using a Qubit 4 fluorometer (Thermo Fisher) and NanoDrop spectrophotometer (Thermo Fisher). Further, RNA was treated with DNAse (DNAse Max, Qiagen, 15200–50) to remove any DNA contamination. Total DNA and RNA were sent to University of Arizona Genomics Core for library preparation and sequencing on an Illumina Novaseq (150bp x 2). DNA was sequenced from four of the eight sampled timepoints (PD, LD, R1, R2; Figure 1), including three replicates from each site, resulting in a total of 24 metagenomes. RNA was sequenced from the eight sampled timepoints, resulting in a total of 48 metatranscriptomes.

**Figure 1.**
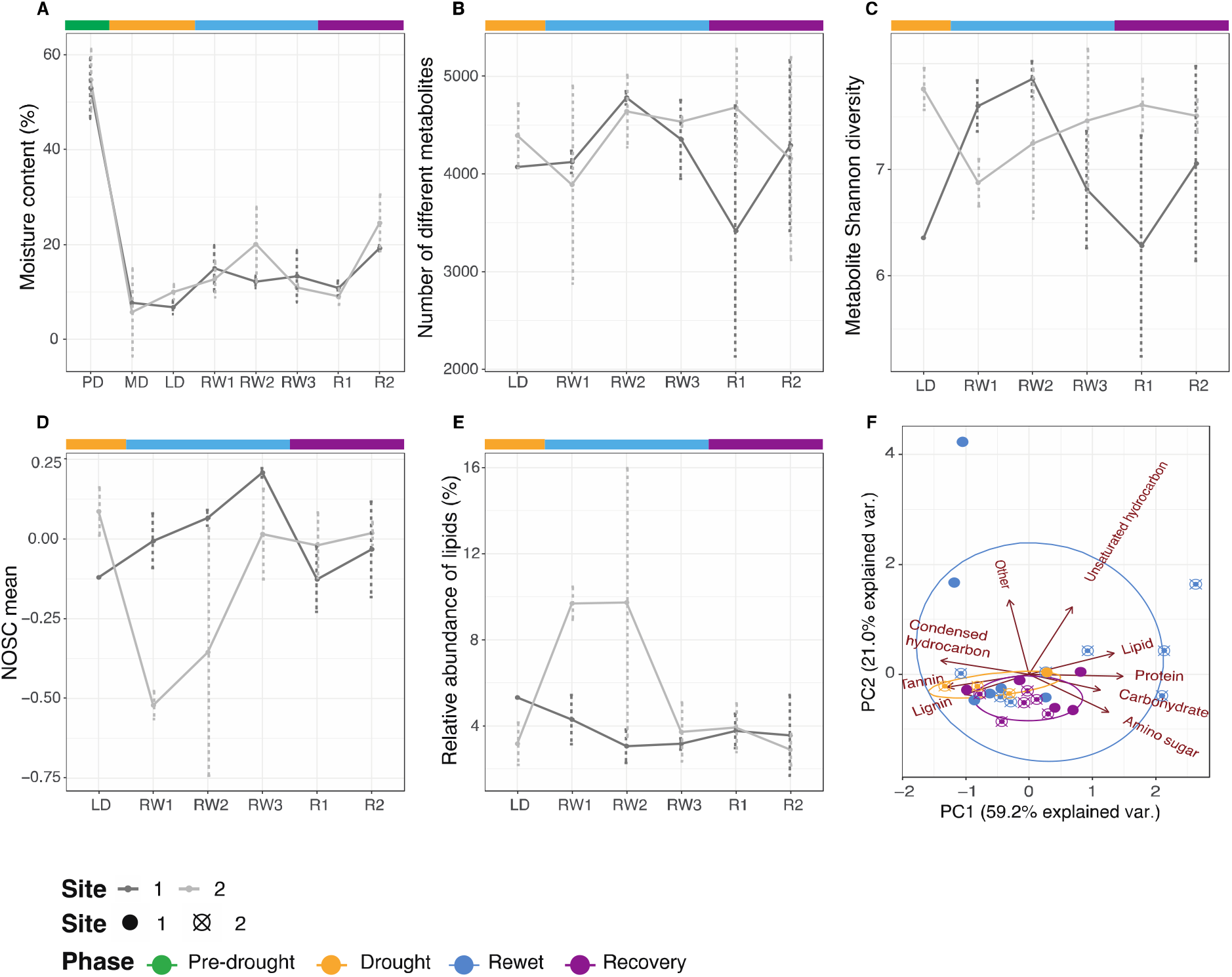
Soil moisture content and metabolite characteristics during drought experiment. Soil moisture content (A), metabolite diversity (B,C), oxidative state of carbon (D) and percentage relative abundance of lipids in soil (E) across timepoints (black line) and between sites (gray lines). (F) Molecular composition of soil metabolome across phases (colors) and sites (shape) with loadings (red arrows and text).

### 2.4. Metagenomic and metatranscriptomic sequence assembly and annotation

Raw reads were processed for quality assessment with FastQC v.0.11.9 (Andrews, 2010), Illumina adapter removal with BBduck v.38.87 (Bushnell, 2022) [ktrim=r k=23 mink=11 hdist=1 tpe tbo] and quality filtering with Trimmomatic v. 0.38 (Bolger et al., 2014) [LEADING:20 TRAILING:20 SLIDINGWINDOW:4:15 MINLEN:50]. An average of 7,327,928 reads per sample were retained. Reads were *de novo* assembled into contigs using MEGAHIT v. 1.1.4 (D. Li et al., 2015), with k-mer length increasing from 21 to 141 in steps of 20. Open reading frames were inferred from contigs using Prodigal v. 2.6 (Hyatt et al., 2010) [-p meta -q -m], and clustered to a 95% similarity 80% coverage using MMseqs2 v.13.45111 (Steinegger and Söding 2017) [--min-seq-id 0.95 --cov-mode 1 -c 0.9 --threads 94] resulting in a total of 29,871,937 non-redundant gene coding sequences. Functional annotation was performed on the non-redundant gene catalog using KofamScan against the KEGG database v.105 (Kanehisa et al., 2017). The abundance of each gene was estimated by mapping to the quality filtered reads using BWA v. 0.7.16 (Li and Durbin 2009) and CoverM v.0.6.1 (Aroney et al., 2024) [-m count --min-read-percent-identity 0.95 --min-read-aligned-length 45], producing a gene count table. We used the resulting KEGG ortholog (KO) number count table to study the abundance of dormancy related genes (sporulation factors, and resuscitation promoting factors; Supplementary Table 3). Counts of the KOs associated with dormancy were normalized by the abundance of species core genes KOs (Supplementary Table 1). Additionally, the cSAP, or community level metabolic niche of heterotrophic bacteria, was inferred from the normalized abundance of KOs associated with sugar and amino and organic acid metabolism. In a nutshell, the cSAP measures the heterotrophic bacteria community’s preference for glycolytic (sugars) or gluconeogenic (amino and organic acids) carbon sources, where a positive value cSAP means that heterotrophic bacteria in that sample show a higher potential for glycolytic metabolism. This measure tells us about bacterial life strategies– glycolytic pathways tend to be favored by fast growing bacteria while gluconeogenic pathways often support bacteria with slower growth – (Gralka et al., 2023). Bacterial taxonomy was annotated from assembled contigs, and a count table of bacterial species was constructed using Metabuli v.1.0.6 (Kim and Steinegger 2024) [--min-score 0.15 --min-sp-score 0.5]. Metabuli uses a combined DNA and six-frame-translated amino acid sequences: it constructs metamers – k-mers that jointly encode DNA and AA information – and compares them to reference databases –GTDB and Refseq – to create the taxonomic classification. Non-bacterial species and bacterial species with a prevalence of less than 10% of the samples, and an abundance of less than 0.00001 were removed for downstream analysis. Finally, bacterial relative abundance was calculated from the count of sequences belonging to a same taxonomic group normalized by the number of reads in the sample.

Paired-end raw RNA sequences were trimmed and filtered using BBduk v.38.87 (Bushnell, 2022) [ktrim=r ordered k=23 mink=11 hdist=1 hdist2=1 minlen=51 minlenfraction=0.33 tbo tpe rcomp=f ftm=5 pratio=G,C plen=20] with a minimum length of 51 and a kmer length of 23. Sortmerna v.4.3.6 (Kopylova et al., 2012) [-fastx] was used to filter out rRNA sequences that matched to the SILVA bacteria/archaea (16S/23S) and eukaryote (18S/28S) databases (Quast et al., 2013). The repair.sh module within BBduk was used to correct paired reads which can get out of order after SortmeRNA. Resulting filtered reads were quality checked with FastQC v.0.11(Andrews, 2010) then assembled using Megahit v.1.2.9 (D. Li et al., 2015) [--presets meta-large] using the “meta-large” presets setting for soil metagenome analysis with high microbial diversity. Contig quality was assessed using Metaquast v.5.2.0 (Mikheenko et al., 2016), and contigs smaller than 200bp were filtered out using SeqKit v.0.3.1.1(Shen et al., 2016).

### 2.5. Viral OTU (vOTU) inference, annotation, and host prediction

Viral inference was performed using an optimized pipeline designed to avoid single tool biases and maximize novel viral genome recovery (Schackart et al., 2023). Briefly, sequences belonging to DNA viruses were inferred from metagenomic assembled contigs using DeepVirFinder v.1.0 (Ren et al., 2020) [-l 1500 -c 8] and VirSorter2 v.2.2.3 (Guo et al., 2021) [-j 4 all --min-length 1500] with a minimum length cutoff of 1500 bp. Inferences from both tools were merged and their quality and completion was assessed using CheckV v1.0.1 tool [end_to_end], and database v.1.4 (Nayfach et al., 2021). Viral inferences of lengths over 5Kb were kept for downstream analysis. While this threshold is lower than some cutoffs reported elsewhere (e.g., 10Kb; Nicolas et al. 2023; Santos-Medellín et al. 2023), they supported recovery of viral sequences for overall community trends from these bulk metagenomes and were subjected to additional filtering criteria. Specifically, only viral sequences detected by Virsorter2 or with a DeepVirFinder score higher than 0.9, that had coverage higher than 5X, and that presented either a viral gene or no host genes detected by CheckV were retained (Schackart et al., 2023). Additionally, we inferred RNA viral sequences from metatranscriptomics using VirSorter2 (Guo et al., 2021) with the same 5 Kb length cutoff. Putative RNA viral contigs with a CheckV score of “Non-determined”, no detected viral genes or at least one cellular gene, and under 5X coverage were discarded. We have chosen an approach that balances scientific thoroughness with the meta-omic practical constraints aiming to capture as much as the viral fraction as possible. VirSorter2 training set includes diverse viral groups (dsDNA, ssDNA, but also and RNA viruses) alongside explicit negative training (Guo et al., 2021). By using VirSorter2 combined with CheckV validation for RNA viral inference (Coclet et al., 2023), we maintain high specificity while maximizing our ability to detect both high and low abundance viral sequences as well as and maintain a consistent approach with the DNA viral inference. Resulting DNA and RNA viral species (vOTUs) were estimated by clustering viral sequences using a 95% identity and 70% completeness cutoff with MMseqs2. Clean metagenomics and metatranscriptomics reads were mapped to viral species to obtain a DNA and RNA vOTU or viral species count table using BWA and CoverM. Finally, DNA viral contigs were mapped against the RNA reads using BWA and CoverM (95% minimum identity and 45 minimum read aligned length) to obtain a measure of the abundance of transcriptionally active DNA viruses. All resulting count tables were manually normalized to reads per kilobase per million (RPKM) using the viral genome sequence length and the number of reads per sample (Coclet et al., 2023). We also normalized all count tables to transcripts per million (TPM) to ensure that our results were not affected by the pitfalls of the normalization method (Zhao et al., 2020).

Potential host taxonomy was predicted from DNA viral sequences using the iPHoP v1.3.3 suite (Shang et al. 2021). In addition, life history classification of viruses in temperate vs. virulent viruses was performed using PhaTYP v.3 (Shang et al., 2023). Finally, we leveraged a list of dormancy-inducing sigma-factor genes previously found in viral genomes (Schwartz et al., 2022). We recovered their sequences from the VOG gene database (Grazziotin et al., 2016) and mapped the sigma-factor gene sequences to our clean RNA reads to estimate their expression levels using BWA and CoverM and posterior RPKM normalization. A schematic overview of the bioinformatics pipelines used in this work, together with all the scripts used to perform the data processing can be found at https://github.com/merytouceda/B2Wald-drought-rewet-virus.

### 2.6. Statistical analyses

We used a linear model to evaluate the differences in richness and abundance of viral species in each phase and site as well as the interaction between these variables. Non-metric dimensional scaling ordination (NMDS) and PERMANOVA test were used to visualize and evaluate, respectively, community composition differences (Bray-Curtis) between phase, timepoint, and site on using the *vegan* package v.2.6-4 (Oksanen et al., 2020). Changes in molecular class composition were visualized using principal component analysis (PCA) and tested also with PERMANOVA. Calculations of Sorensen dissimilarity, and its components, nestedness and turnover on the spatial and temporal scale were calculated using the *betapart* package v.1.6. (Baselga and Orme 2012), and the differences were tested with a Wilcoxon rank sum test with continuity correction to evaluate how viral communities vary across space and time and determine whether the differences (Euclidian) in community composition are caused by richness differences (higher nestedness), or by species replacement (higher turnover). Finally, the same interaction model between phase and site was used in differential abundance analyses of the DNA viral hosts using MaAsLin2 v.1.12 (Mallick et al., 2021). All statistical analyses and visualizations were implemented in R v.4.2.2 (R Core Development Team, 2015) and can be found at https://github.com/merytouceda/B2Wald-drought-rewet-virus.

## 3. Results

### 3.1. Soil moisture content and soil metabolites are determined by site and in some cases experimental phase differences

Soil moisture content fluctuated significantly across experiment phases (pre-drought, drought, re-wet, and recovery). In both sites, soil moisture content decreased dramatically during drought, started to recover upon rewet and continued to increase towards the end of the recovery phase, although not achieving pre-drought (PD) levels (phase: F = 177.10, P < 0.001; site: F = 0.84, P = 0.369; Figure 1A). Although water content did not recover during rewet, water potential measurements at slightly more shallow layers (Supplemental Figure 1B) reveal that the water stress was significantly relieved during the rewet and achieved recovery by the end of this period. Soil metabolite composition and diversity showed complex responses to drought and rewetting that varied by site. While the total number of metabolic compounds showed no significant variation across phases and sites (Table 1), both sites exhibited increased metabolite numbers during rewetting followed by site-specific patterns during recovery (Figure 1B). Metabolite Shannon diversity revealed non-significant but contrasting site-specific trends (Table 1; Figure 1C). Site 1 showed increased diversity during rewetting followed by a decrease during recovery, while Site 2 displayed the opposite pattern. The nominal oxidation state of carbon (NOSC) reflects the average oxidation state of carbon atoms in a molecule and closely associated with thermodynamic favorability of degradation, especially for microbial respiration, and thus serves as an indicator of organic matter quality for decomposition (Wilson and Tfaily 2018). The NOSC also showed non-significant but contrasting site-specific responses (Table 1; Figure 1D). NOSC peaked 48 hours into rewetting (RW2) in Site 1 with a steady increase, while Site 2 showed a rapid decrease during rewetting. Both sites returned to pre-drought conditions during recovery. In contrast, the relative abundance of lipids, which reflects the composition of the soluble lipid pool that is most likely bioavailable or mobile in the environment, dropped during rewet in Site 1, and increased during rewet in Site 2, both sites showing similar levels at the end of the rewet and during recovery, and again showing no significant changes (Table 1; Figure 1E). Finally, the effect of the experimental phase on soil molecular composition depended on the site (Table 1; Figure 1F).

**Table 1.**
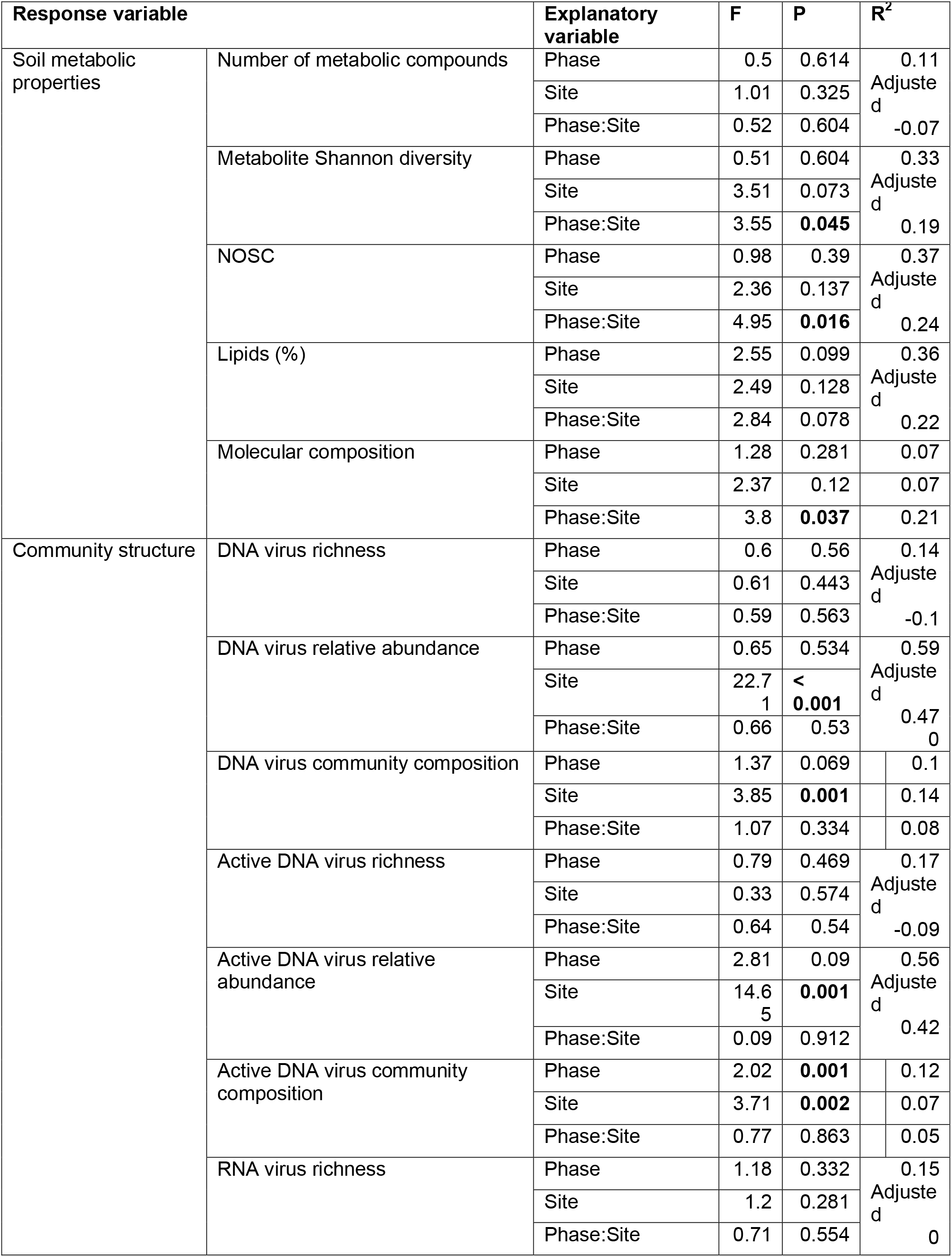
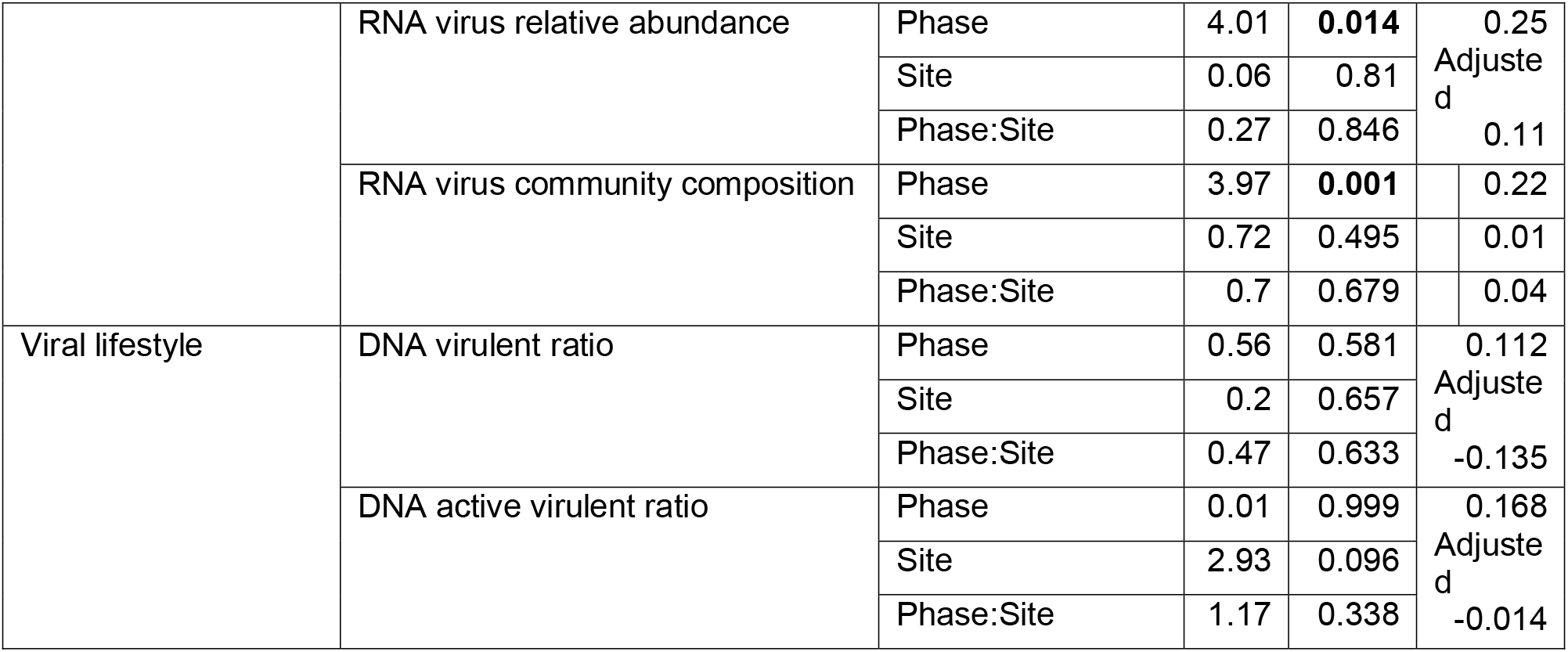
Statistical results summarizing how soil metabolic properties, viral community structure and viral lifestyle vary based on experimental phase and site. A linear model accounting for the interaction between these variables was implemented using ANOVA for numeric variables, and PERMANOVA for community composition. Relative abundance measures are normalized to RPKM.

### 3.2. Community structure of DNA and RNA viruses mostly differs among sites, while the abundance of active DNA viruses fluctuates with the experimental phase

We recovered 6,452 DNA and 819 RNA viral species from the metagenomic and metatranscriptomic assemblies, respectively. Species richness did not differ significantly (Table 1) between experiment phase (pre-drought, drought, recovery) for either DNA viruses (Figure 2A), transcriptionally active DNA viruses, (Figure 2B), or RNA viruses (Figure 2C). However, DNA and RNA viral richness followed a similar non-significant pattern, reaching the highest levels at the start of the recovery phase (R1) and lowest during the drought (MD or LD; Figure 2). Similarly, although the relative abundance (in RPKM) of DNA, or transcriptionally active DNA, did not differ significantly between phases (Table 1; Figure 2D,E), we also observed similar patterns in the relative abundance of transcriptionally active DNA and RNA viruses, which did differ significantly between phases (Table 1; Figure 2E,F). Interestingly, we noticed that the relative abundance of DNA and transcriptionally active DNA viruses were especially influenced by site (Table 1; Figure 2D,E). The same overall patterns, which slight changes in site-specific patterns, were present when using TPM, instead of RPKM, normalized counts (Supplementary Figure 2 and Supplementary Table 4).

**Figure 2.**
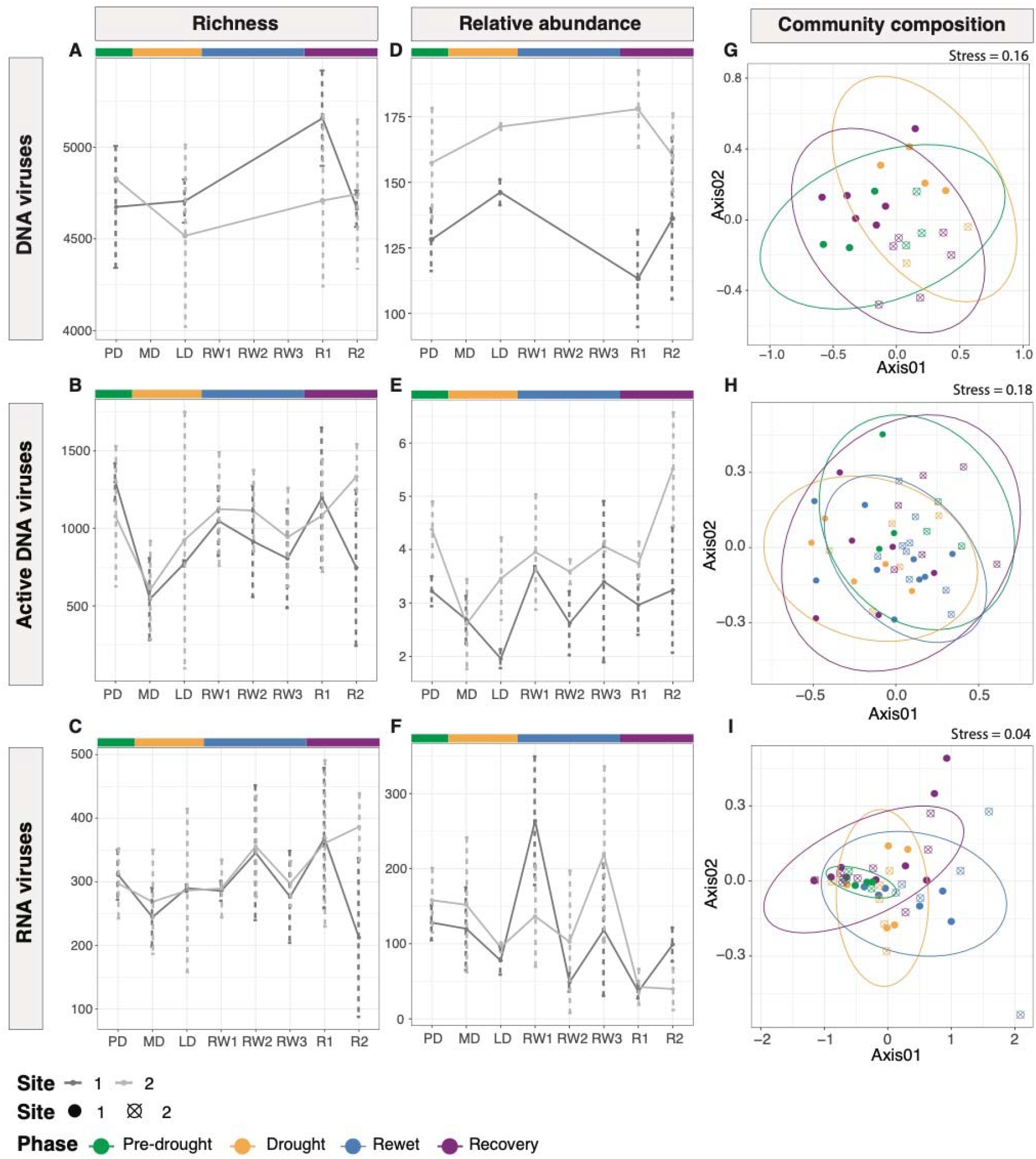
Changes in the diversity, RPKM-normalized abundance, and composition of DNA viruses, active DNA viruses, and RNA viruses during drought experiment. DNA, transcriptionally active DNA, and RNA species richness (A-C) and relative abundance in reads per kilobase per million (RPKM; D-F) across timepoints (black line) and between sites (gray lines). Viral community composition across phases (colors) and sites (shape). Ellipses denote experiment phase 95% confidence intervals.

We observed high overlap in viral species among all studied phases (Supplementary Figure 3). Nevertheless, community composition was different between phases both for transcriptionally active DNA viruses and RNA viruses (Table 1; Figure 2H,I). Particularly, DNA viruses that infect bacteria of the families Pseudomonaceae (phylum Proteobacteria) and B64-G9 (phylum Myxococotta) were transcriptionally active in the recovery phase compared to pre-drought communities, but no differences were observed in the other phases. Furthermore, we found that the composition of DNA viral communities and transcriptionally active DNA viral communities was strongly determined by site (Table 1; Figure 2G,H). In fact, DNA viral communities showed higher levels of spatial Sorensen dissimilarity (W = 3560, P < 0.001; Supplementary Figure 4A) and spatial turnover (W = 3552, P < 0.001; Supplementary Figure 4C), while nestedness did not change significantly in the temporal scale (W = 2458, P = 0.753; Supplementary Figure 4B).

### 3.3. Bacterial community structure was determined by site and phase, with certain phyla preferring different phases

Since they are the main host to DNA viruses we studied the structure of bacterial communities to evaluate their association with viral community structure. Similarly to DNA viruses, we observed no differences in bacterial richness (F = 3.35, P = 0.061; Supplementary Figure 5A) or relative abundance (F = 0.06, P = 0.969; Supplementary Figure 5B) between phases, nor sites (richness: F = 0.20, P = 0.659; relative abundance: F = 2.54, P = 0.131). Bacterial richness and relative abundance were positively correlated with DNA viral richness, but not with DNA viral relative abundance or transcriptionally active DNA viral richness and relative abundance (Supplementary Figure 6). Furthermore, bacterial richness had a negative association with soil moisture content (Supplementary Figure 6). We found that bacterial community composition was determined by the interaction between site and phase (R_2_ = 0.15, P = 0.045; Supplementary Figure 5C). Moreover, we found certain bacterial phyla to be differentially abundant between phases. Specifically, the phylum Actinomycetota were less abundant in the pre-drought and recovery phases compared to the drought, while the relative abundance of Pseudomonadota was higher in the recovery phase compared to the drought (Supplementary Figure 5D).

### 3.4. DNA viruses were predominantly virulent, while bacteria showed a preference for organic acid-based metabolism

We predicted the viral lifestyle (temperate vs. virulent) of 83% of DNA viral genomes— expressed as ratios of relative abundance and expression of virulent vs. temperate viruses. Neither of these were significantly determined by phase or site (Table 1, Supplementary Figure 7).

To evaluate the phenotypic shifts that could be induced in bacterial communities due to resource availability, we calculated the community weighted bacterial preference for sugar vs. aminoacidic compounds from the normalized abundance of genes associated with their metabolism (cSAP). We found levels of cSAP to be consistently negative, suggesting that these bacterial communities favor organic acid catabolism. Again, we observed that the preference for sugar or amino and organic acid carbon sources did not depend on the phase or site (phase: F = 0.86, P = 0.441; site: F = 0.28, P = 0.603; Supplementary Figure 5E). Finally, our search of sigma factors related to host dormancy in soil viruses did not yield any results, as we did not find any sigma factors encoded in viral genomes (data not shown). Similarly, no bacterial dormancy related gene, either sporulation factor or resuscitation factor was differentially abundant in a specific phase (Supplementary Figure 5F, G).

### 3.5. The abundance and transcriptional activity of DNA viruses was positively associated with metabolite diversity and the mean NOSC

To illustrate the potential relationships between soil viruses and ecosystem processes we analyzed the associations between viral and metabolomic parameters. The relative abundance of DNA viruses was positively correlated with the Shannon diversity of metabolomic compounds (Supplementary Table 5; Figure 3A), and the mean nominal oxidation state of carbon (NOSC; Supplementary Table 5; Figure 3B), and showed a negative trend association, although not statistically significant, with the percentage of lipids (Supplementary Table 5; Figure 3C). We found a non-significant but positive tendency in the association between the relative abundance of transcriptionally active DNA viruses and metabolite Shannon diversity (Supplementary Table 5; Figure 3E), but no correlation with the mean NOSC (Supplementary Table 5; Figure 3F), and the percentage of lipids (Supplementary Table 4; Figure 3G). We found no relationship between the relative abundance of DNA viruses and the moisture content (Supplementary Table 5; Figure 3D), or the relative abundance of transcriptionally active DNA viruses and the moisture (Supplementary Table 5; Figure 3H). Finally, we found no significant (P < 0.05) or strong (Pearson’s correlation: r > [0.4]) correlations between RNA viral abundance and metabolite Shannon diversity, NOSC, percentage of lipids, or moisture content (Supplementary Table 5; Figure 3I-L). All associations between viruses and metabolite soil characteristics were highly dependent on site (Supplementary Figure 8). Additionally, correlations maintained the direction but lost the statistical significance when using abundance in TPM instead of RPKM (Supplementary Figure 9), and again when using TPM, we observed differences in associations based on site (Supplementary Figure 10).

**Figure 3.**
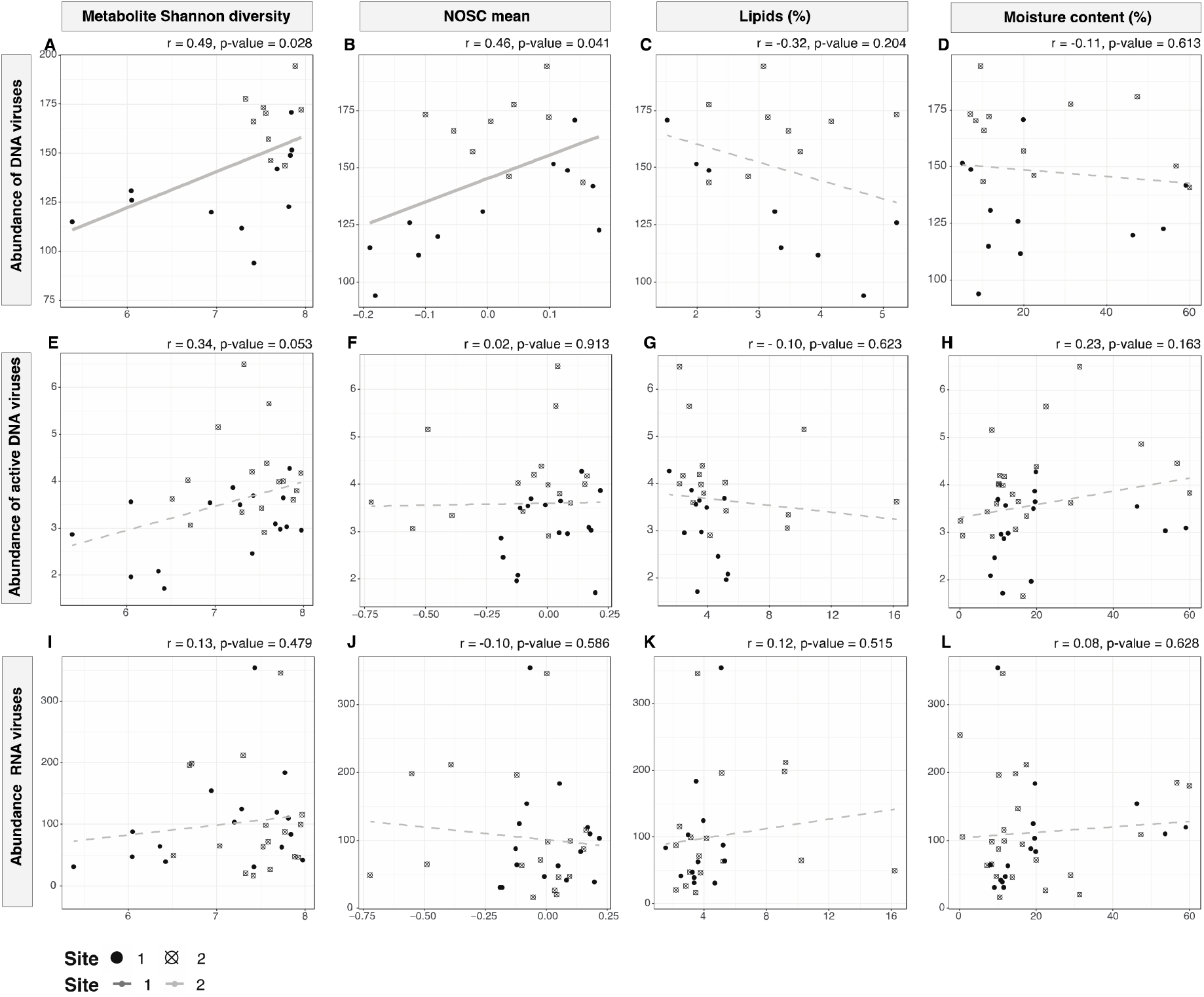
Site-consolidated associations between DNA, transcriptionally active DNA, and RNA viral RPKM-normalized abundance, metabolite diversity, and carbon composition measures. Lines represent the estimated linear association between variables (dark gray = site 1, light gray = site 2). Pearson correlation results are placed on the top right of each plot. Significant correlations are marked by a solid line. Point shape is assigned based on site identity (solid point = site 1, point with crossing lines = site 2).

## 4. Discussion

Understanding the responses of soil microbial communities to changes in ecosystem dynamics is increasingly important in the face of climate change (Cavicchioli et al., 2019; Classen et al., 2015). Soil viruses have been identified as key players in the cycling of carbon marine ecosystems (Fuhrman, 1999; Suttle, 2007), while their role in carbon and organic matter mobilization in soils is increasingly recognized (Trubl et al., 2018). Tropical ecosystems are projected to experience a higher number and intensity of droughts as a result of climate warming, with potential consequences on soil biogeochemical cycling and microbial function (Reed et al., 2020; Cusack et al., 2024). In this study, we combined meta-omics data to investigate the response of DNA and RNA viruses to drought, and their relationship with soil organic carbon in an artificial tropical rainforest.

Viral life in the soil is highly dependent on moisture. In dry soils, viruses are constrained to shrinking water films and reduced contact with bacterial cells, which limits their reproduction. When soil is rewetted, water fills the pores, which could enable viruses to move freely, encounter and infect new bacterial populations, and begin fresh cycles of lytic replication (Williamson et al., 2017). Although we did not observe significant changes in the diversity and relative abundance of DNA viruses, transcriptionally active DNA viruses and RNA viruses, as a response to changes in soil moisture content, we did observe a non-significant trend of decrease in diversity and relative abundance as soil moisture content dropped during drought, followed by peaks during the rewet and recovery phases in both sites for active DNA viruses and RNA viruses. It is worth noting that the soil moisture content remained under 25% after drought, which could explain the lack of strong patterns in soil viral diversity and relative abundance. Additionally, we found no association between viral relative abundance and soil moisture, indicating that viral dynamics likely depend on unexplored environmental or biological factors beyond the parameters of this study. Along those lines, we observed that the richness, relative abundance, and community composition of overall DNA viruses were determined by site rather than changes in soil moisture content. These results highlight the intricate ecological dynamics of soil ecosystems, where complex interactions can magnify subtle environmental changes. For instance, a rewetting event can trigger a myriad of responses through the redistribution of accumulated nutrients within the soil’s heterogeneous structure (Kuzyakov and Blagodatskaya 2015). In fact, DNA viruses primarily infect bacteria, whose community structure is determined by multiple abiotic factors and soil chemistry– particularly pH (Fierer and Jackson 2006)–potentially obscuring the influence of the changes in soil moisture. Interestingly, studies have reported a similarly important role of spatial heterogeneity in the structuring of the microbial function in the same system (Honeker et al., 2023) and of viral communities in natural systems (Barnett and Shade 2024). This is a relevant issue to be considered when designing time-scale studies on soil viruses, since strong spatial heterogeneity poses an added difficulty for the prediction of soil viral response to environmental changes.

Contrary to our expectation, we did not observe a return to pre-drought diversity, relative abundance, or community composition of viral communities when precipitation dynamics were recovered. This could be caused by the moisture content not having returned to pre-drought levels entirely or could imply that viral communities were irreversibly departed from their pre-drought compositions after drought disturbance, and thus might not be resilient over the timescale evaluated. This contrasts with previous studies on the effects of drought on prokaryotic communities in rainforest, which show communities that diverge during drought but return to pre-drought levels soon after the re-introduction of precipitation (L. Li et al., 2021). Perhaps a continuation of the experiment would have seen the viral community composition return to pre-drought conditions as soil moisture reached pre-drought levels. However, it is still possible for the variability among small scale soil habitats, and among different types of tropical forests to lead to discrepancies in their responses to disturbances like drought (Tripathi et al., 2016).

Altogether, our results suggest that spatial patterns and heterogeneity are important controllers of viral responses. Furthermore, viral communities of the Biosphere 2 artificial rainforest showed a reduced overall response compared to what has been reported in other ecosystems. However, a majority of these studies have been done on soils of arid or semiarid regions (Santos-Medellín et al., 2023), soils of thawing permafrost (Emerson et al., 2018), or watersheds (Coclet et al., 2023), whose ecosystem dynamics include natural periods of drought and rewet. Even among studies centered in similar ecosystems, the response of soil viral communities to changes in moisture is not consistent, with reports of both increase (Santos-Medellín et al., 2023) and decrease (Nicolas et al., 2023) in viral richness after dry arid or semi-arid soils wet up.

Another possible explanation for the lack of viral response we observed is that our study focused on metagenome-inferred viral sequences, which are less reactive than viromes obtained through first retrieving viral particles and posterior sequencing (Santos-Medellin et al., 2021). Although these limitations are not unusual in current soil virology, and even unavoidable in certain studies, like those using pre-existing datasets, we believe that new experimental designs should adopt a combination of metagenome-inferred viral sequences and viromes to differentiate between prophage (i.e. a virus integrated in the bacterial host genome) and extracellular viral populations. Soil viral research is still an emerging scientific discipline whose advance is largely tied to in-silico techniques that are being iteratively refined to develop more robust analytical standards (Roux et al., 2019). Here, we found that a relatively low inference threshold was required to recover DNA viral sequences from metagenomes compared to other studies (Nicolas et al. 2023; Santos-Medellín et al. 2023). Encouragingly, this method revealed consistent and significant patterns in DNA viral abundance and associations (e.g., with soil carbon).

Previous studies from the B2 WALD campaign have documented drought-induced changes in carbon mobilization (Werner et al., 2021; Honeker et al., 2023; Huang et al., 2024). Given established experimental (Braga et al., 2020; Albright et al., 2022;) and observational (Trubl et al. 2018; Van Goethem et al. 2019; Barnett and Buckley 2023) evidence linking viral activity to nutrient mobilization in soils, we investigated relationships between viruses and organic carbon molecular composition. Our results revealed positive correlations between active DNA viral abundance and metabolite diversity in one of the sites (Site 1), suggesting DNA viral replication may influence nutrient mobilization through cell lysis (Heinrichs et al., 2020). Furthermore, higher DNA virus abundance correlated in one of the sites with more oxidized carbon states and showed a non-significant negative association with the relative abundance of stable carbon molecules like lipids. These patterns could be explained by bacterial cell lysis releasing endoenzymes that trigger extracellular oxidative metabolism (Maire et al., 2013) and providing labile material for surviving microorganisms, thereby enhancing microbial respiration of complex molecules (Kuzyakov and Blagodatskaya 2015). The absence of correlations between viral and bacterial abundance suggests these results are not simply due to increased prophage abundance. While these correlations do not demonstrate causation, which would require viral addition experiments (Blazanin and Turner 2021; Albright et al. 2022), they highlight viruses’ potential contribution to nutrient mobilization and carbon cycling, emphasizing the importance of understanding viral responses to environmental change. The differences we found in viral community structure between sites evidences the importance of unidentified environmental conditions on soil viral function (Zimmerman et al., 2024). In our system, DNA viruses exhibited a higher sensitivity to spatial heterogeneity, while RNA viral communities and secondary changes in DNA viral populations responded dynamically to different experimental phases. This suggests that microlevel changes in soil moisture and other environmental conditions could potentially influence viral community composition. Additionally, the association between DNA viral abundance and metabolite diversity provides insights into the potential mechanisms by which changes in soil viral structure might impact broader soil processes (Nicolas et al., 2023). The observed patterns raise important questions about the potential ecological implications of viral community dynamics, particularly in the context of prolonged drought in tropical rainforests. Future research is needed to fully understand how viral community changes could potentially affect carbon dynamics and microbial biomass turnover(Zimmerman et al., 2024).

Along those lines, and despite those associations, we found no changes in the abundance or activity of viruses with different life strategies (i.e. virulent vs. temperate). Soil viruses are one of the most abundant biological entities known, but their study has only recently been made possible through metagenomics and advances in viral sequence inference tools (Schackart et al., 2023). Viral lifestyle annotation provides a framework to make sense of this large amount of high complexity data (Barberán et al., 2012). However, current tools for the characterization of viruses rely on functional annotations of viruses, which are still limited (i.e. we did not find any known sigma-factor genes in our viral genomes). Additionally, functional annotations can speak of the genetic potential of a virus to carry a lifestyle but serve as flawed predictions of their behavior. For example, temperate viruses can start lytic reproduction life cycles when conditions are optimal, or viruses with virulent lifestyles can fail to contact a host and therefore not produce infection (Chevallereau et al., 2022). Ultimately, our study reveals the challenges in predicting viral behavior through genetic annotations, highlighting the need for improved approaches, such as coverage-based analysis of temperate phage activity (Kieft and Anantharaman, 2022), to understand viral ecological interactions (Camargo et al. 2023; Liu et al. 2023; Roux et al. 2023; Hou et al. 2024).

## 5. Conclusion

In this meta-omics study of the viral community dynamics during a 60-day drought in an artificial tropical rainforest, we demonstrated that although soil viral communities respond to changes in the environment, their response can be highly dependent on small-scale soil heterogeneity, and therefore hard to predict at the ecosystem level. Moreover, we discussed the impacts that changes in viral abundance can produce in soil nutrient, and specifically, carbon cycling; higher DNA viral abundances being associated with higher metabolite diversity. Since climate change will bring dramatic changes in ecosystems dynamics across the globe, it is important that we understand the effects of these changes in microbial communities with an underexplored role in biogeochemical cycling such as soil viruses.

## Data availability

The metagenomics and metatranscriptomics raw sequence data are publicly available through Genbank SRA under the following BioProject IDs: PRJNA980752– PRJNA980834. Processed data products for the study of viruses and metabolites are available at https://zenodo.org/records/14720854. All code for data analysis and figure creation can be found at https://github.com/merytouceda/B2Wald-drought-rewet-virus

## Acknowledgements

We thank Michaela Dippold for acquiring some of the soil samples.

