## Supplementary Information for "Soil viral response to drought disturbance and ecosystem recovery in an artificial rainforest"

**Supplementary Methods**

**Harnessing metagenomics to unveil the underrepresented fraction of the soil microbiome**

**Soil description**

The soil in the Biosphere 2 Tropical Rainforest (TRF) biome was artificially constructed to replicate the functional characteristics of natural tropical rainforest soils while providing a stable substrate for long-term plant growth. The soil system consists of two main layers, a *subsoil* that that varies in thickness, and it is composed of a rocky, pebbly sandy loam extracted from a local quarry and selected to improve water percolation.

The *topsoil* is a from a silt loam from a local desert grassland with organic amendments made of forest mulch, alfalfa, cotton gin trash and manure. Although, the TRF has a uniform composition, the total soil depth varies within the different TRF areas has a range from 1.8 to 2.9-m for Site 1 and from 2.5 to 3.3m for Site 2.

Overall, the soil texture is classified as sandy clay loam (18-24% Clay, 36 - 46% Silt, 36-39% Sand, 15-20% Gravel).

The latest soil physico-chemical description of the different areas can be found as Table 1 in Pugliese et.al. (2023).

[Pugliese, G., Ingrisch, J., Meredith, L. K., Pfannerstill, E. Y., Klüpfel, T., Meeran, K., ... & Williams, J. (2023). Effects of drought and recovery on soil volatile organic compound fluxes in an experimental rainforest. nature communications, 14(1), 5064.](https://www.nature.com/articles/s41467-023-40661-8)

**Supplementary Figures**

**Supplementary Figure 1. Experimental design and soil water matric potential. A.** Black dots denote sampling events with timepoint abbreviations: pre-drought (PD), mid-drought (MD), late drought (LD), rewet (RW), and recovery (R). **B.** Soil water matric potential across timepoints and experiment phases as the average of soil water potential sensors buried at 5 and 10 cm (output WP kPa and T °C 693 TEROS 21, Meter Group, Pullman, WA, USA)(Werner et al., 2021). **C.** Schematic of the 0.19 ha of artificial tropical rainforest where the experiment was conducted. Yellow shapes represent the two sites where soil samples were taken. The maximum distance between the 2 outer most points of the areas are approximately 20 meters and the minimum distance was 1m.

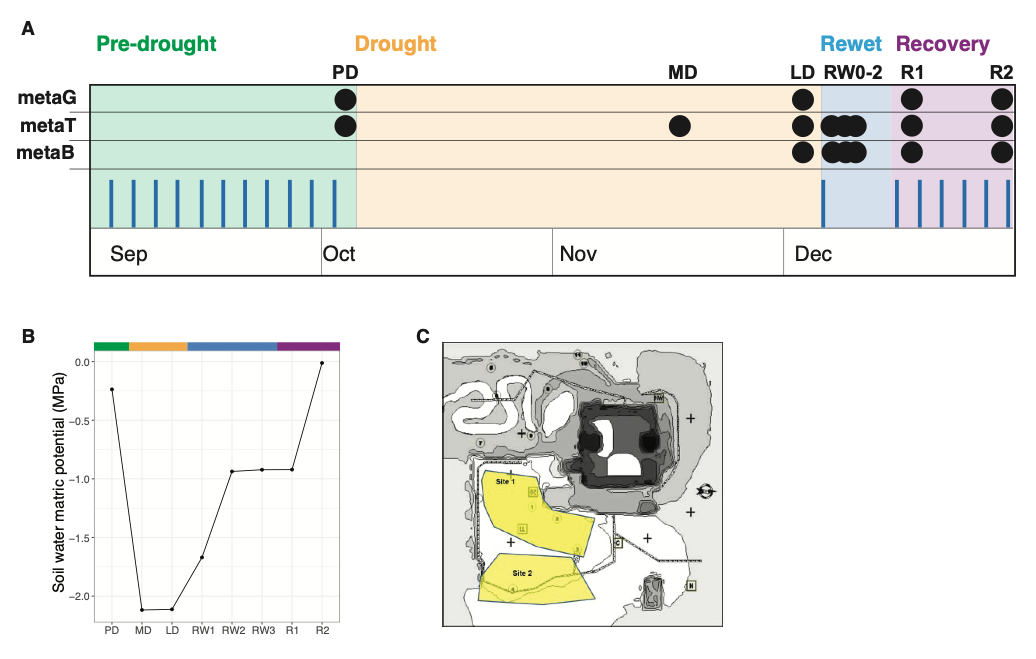

**Supplementary Figure 2. Changes in the diversity, abundance, and composition of DNA viruses, active DNA viruses, and RNA viruses during drought experiment.** DNA, transcriptionally active DNA, and RNA species richness (A-C) and abundance in transcripts per million (TPM) (D-F) across timepoints (black line) and between sites (gray lines). Viral community composition across phases (colors) and sites (shape). Ellipses denote experiment phase groupings.

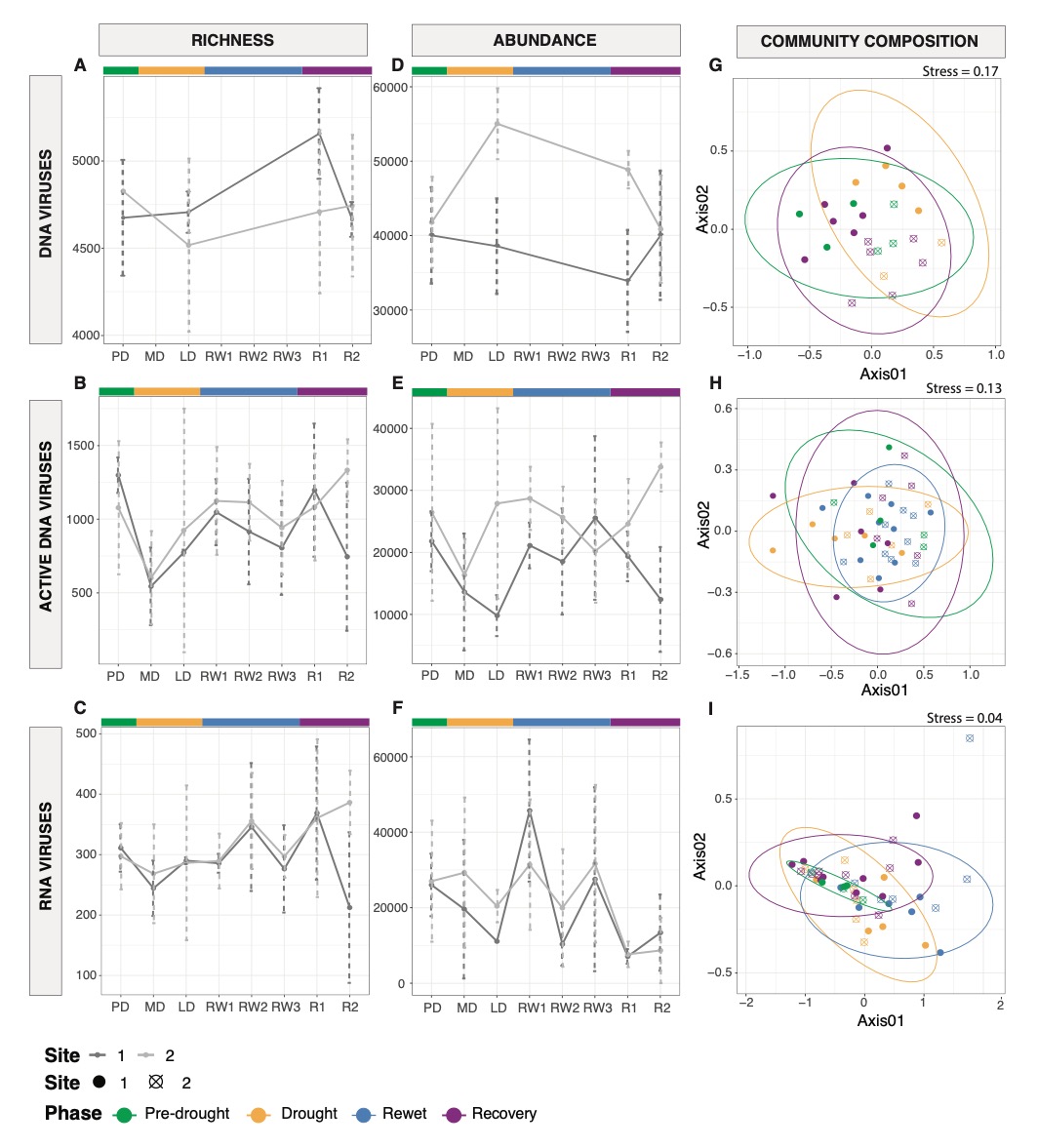

**Supplementary Figure 3. Species overlap between phases.** Upset plots of DNA viral species, active DNA viral species**,** and RNA viral species (top to bottom) separated by site (site 1 = left, site 2 = right). Columns represent the number of shared species among the soils of each timepoint (rows with points). Colors denote phase of experiment: green for pre-drought, yellow for drought, blue for rewet, purple for recovery.

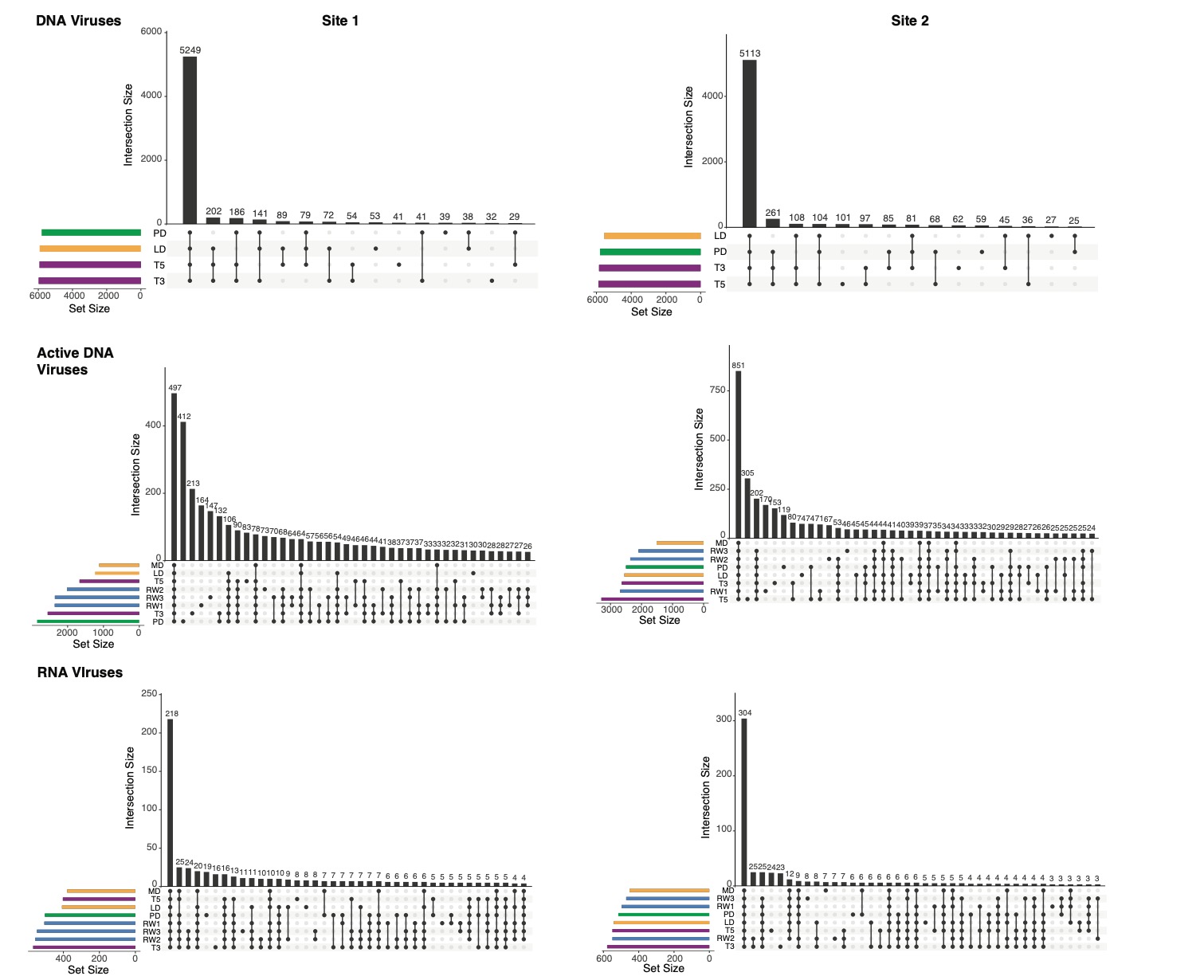

**Supplementary Figure 4. Beta-diversity components of viral communities.** Sorensen dissimilarity (top row) and its components: nestedness (middle row) and turnover (bottom row) at the spatial (between sites) and temporal (between timepoints) scale for DNA viruses, transcriptionally active DNA viruses, and RNA viruses. Significant results denoted by red font.

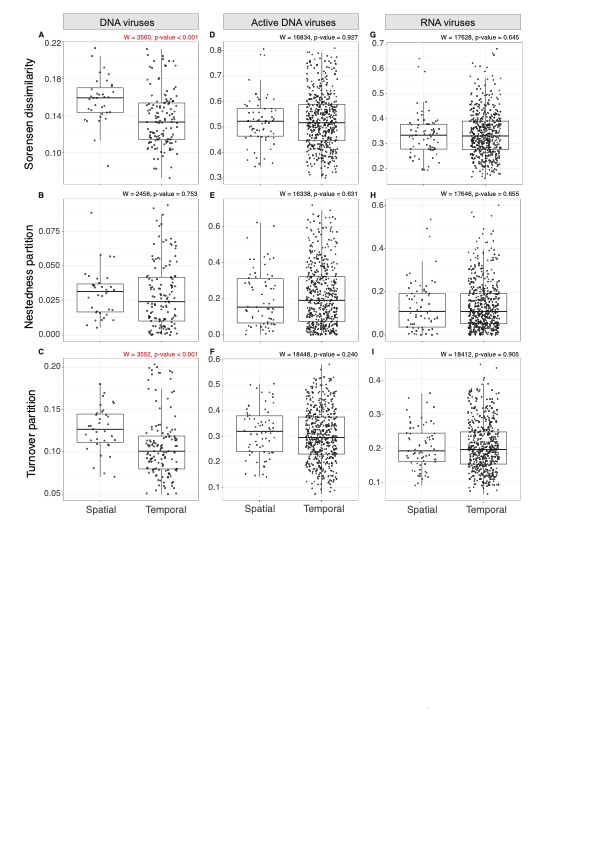

**Supplementary Figure 5. Bacterial community structure and functional traits.** Bacterial species richness (A) and RPKM-normalized abundance (B) at different timepoints and sites (gray scale). Bacterial community composition (C) and phase based differentially abundant major phyla (D). Community weighted sugar-acid preference in bacteria (E) and dormancy genes (sporulation and resuscitation) found in community (F-G).

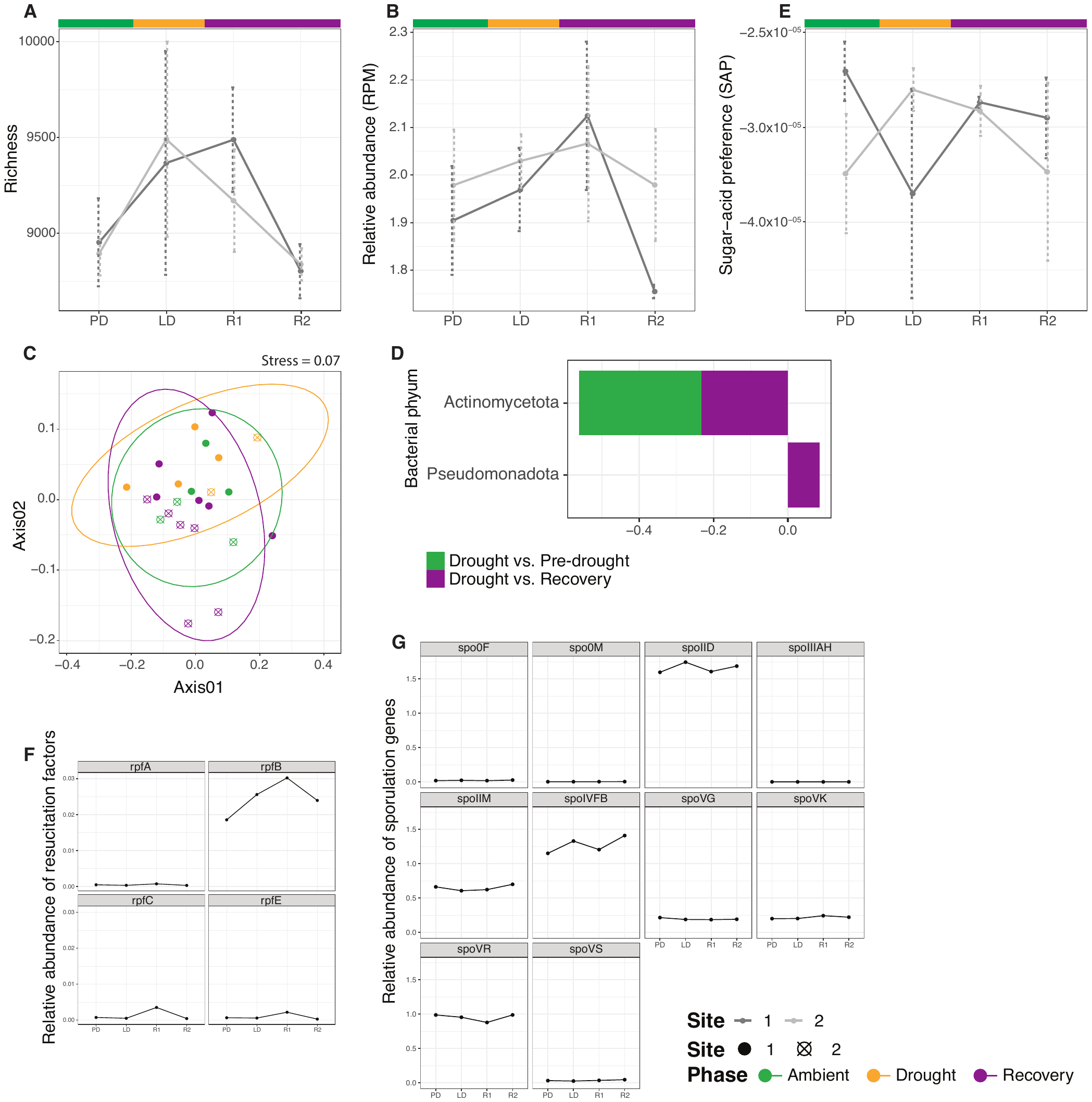

**Supplementary Figure 6. Associations between bacterial community structure, viral community structure, and moisture.** Correlations (Pearson) between bacterial richness and RPM-normalized abundance measures and viral richness and RPKM-normalized abundance measures, and soil moisture content.

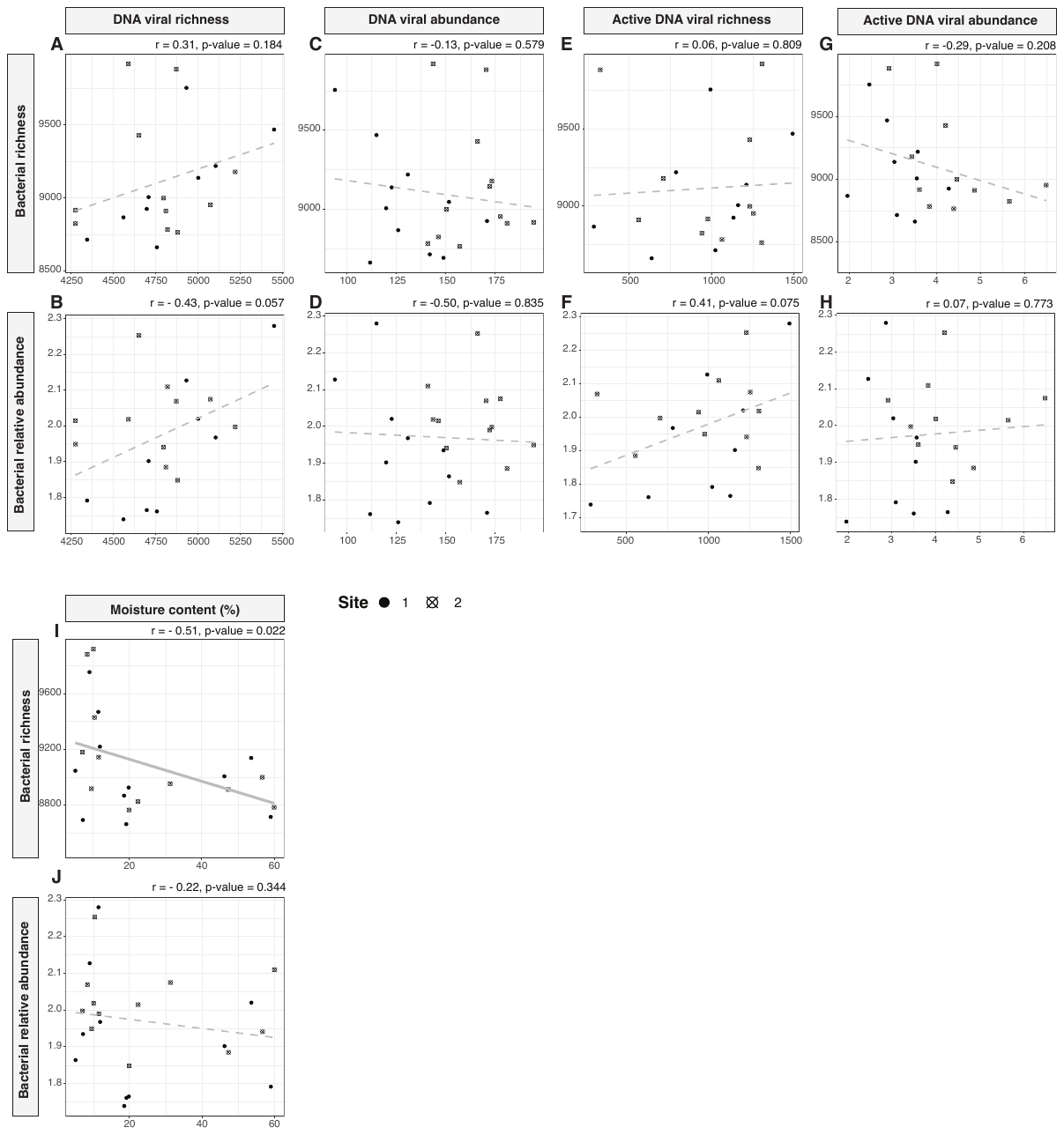

**Supplementary Figure 7. Abundance and expression ratio of virulent vs. temperate viruses.** Abundance ratio (A) and expression ratio (B) of virulent vs. temperate viruses across timepoints (black line) and between sites (gray lines).

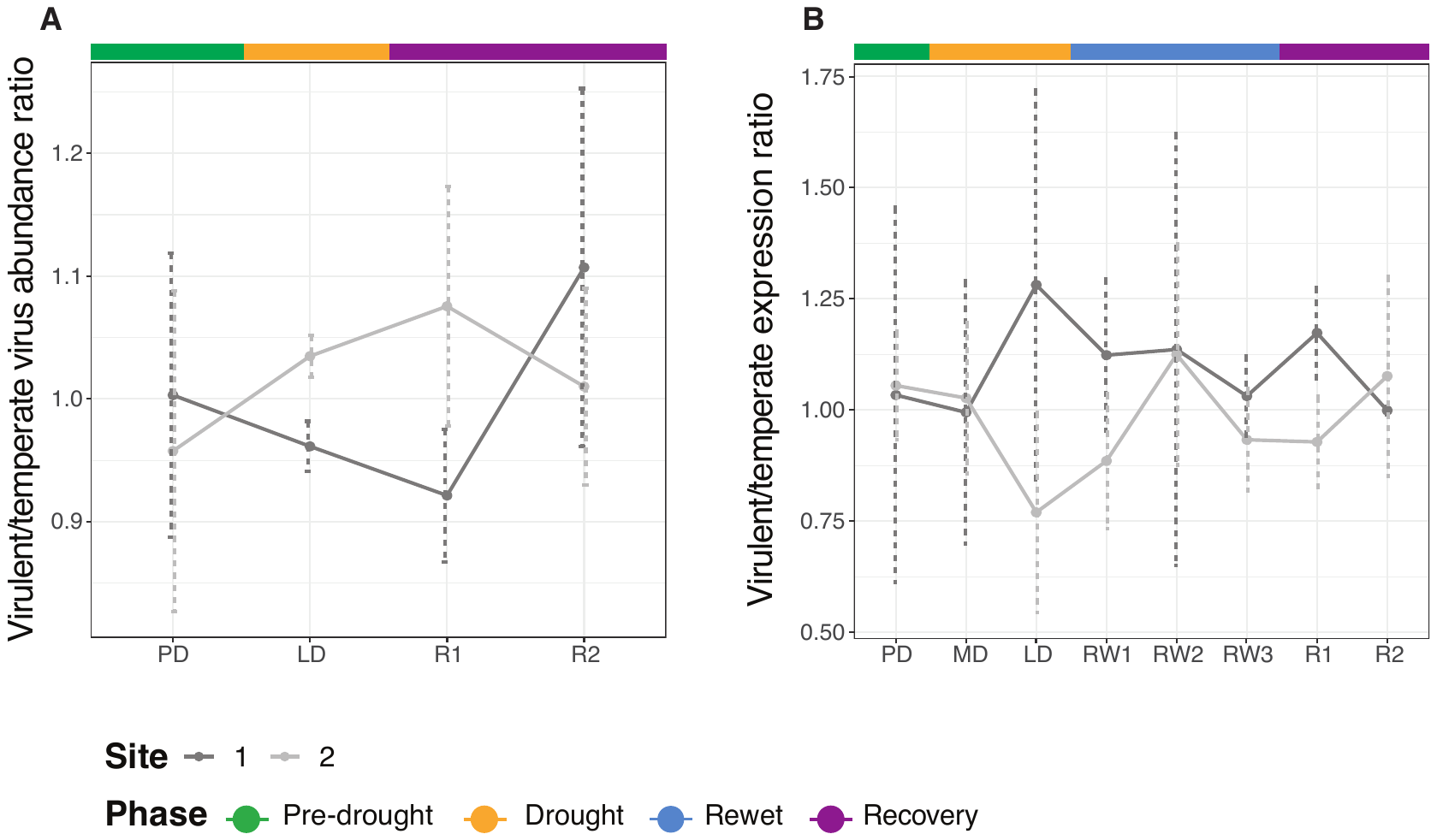

**Supplementary Figure 8. Site-specific associations between DNA, transcriptionally active DNA, and RNA viral RPKM-normalized abundance, metabolite diversity, and carbon composition measures.** Lines represent the estimated linear association between variables (dark gray = site 1, light gray = site 2). Pearson correlation results are placed on the top right of each plot. Point shape is assigned based on site identity (solid point = site 1, point with crossing lines = site 2).

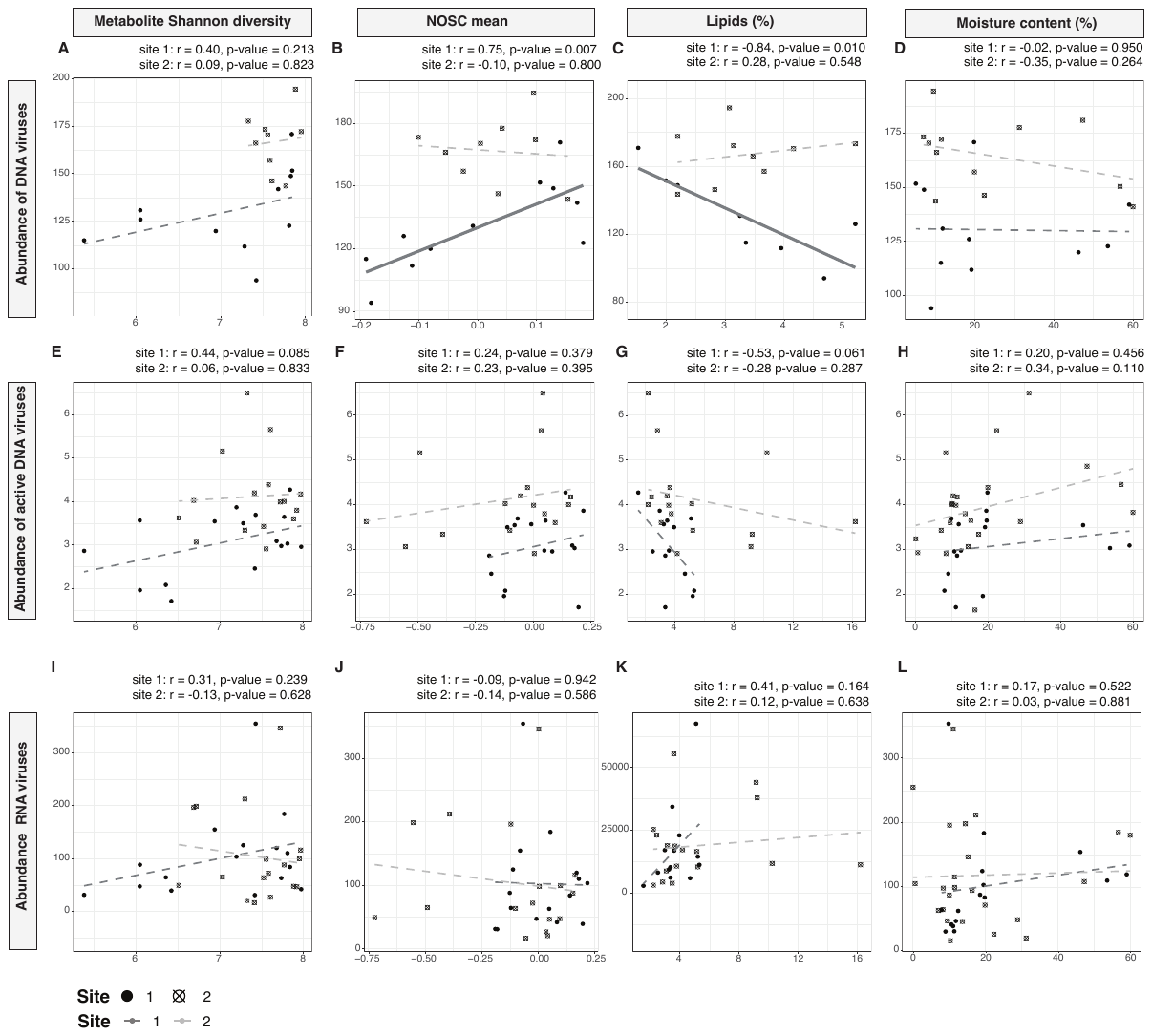

**Supplementary Figure 9. Site-consolidated associations between DNA, transcriptionally active DNA, and RNA viral TPM-normalized abundance, metabolite diversity, and carbon composition measures.** Lines represent the estimated linear association between variables. Pearson correlation results are placed on the top right of each plot. Point shape is assigned based on site identity (solid point = site 1, point with crossing lines = site 2).

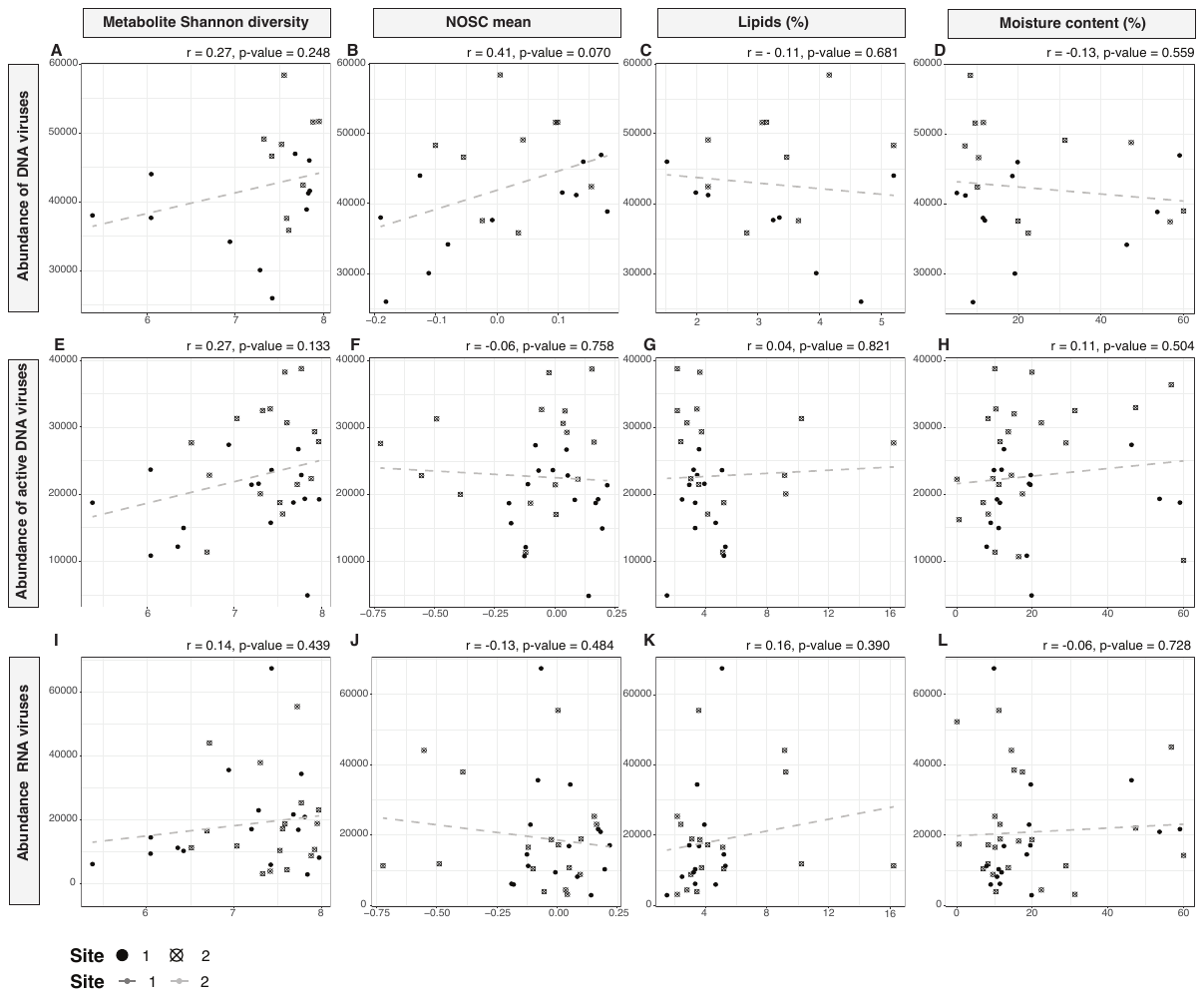

**Supplementary Figure 10. Site-specific associations between DNA, transcriptionally active DNA, and RNA viral TPM-normalized abundance, metabolite diversity, and carbon composition measures.** Lines represent the estimated linear association between variables (dark gray = site 1, light gray = site 2). Pearson correlation results are placed on the top right of each plot. Point shape is assigned based on site identity (solid point = site 1, point with crossing lines = site 2).

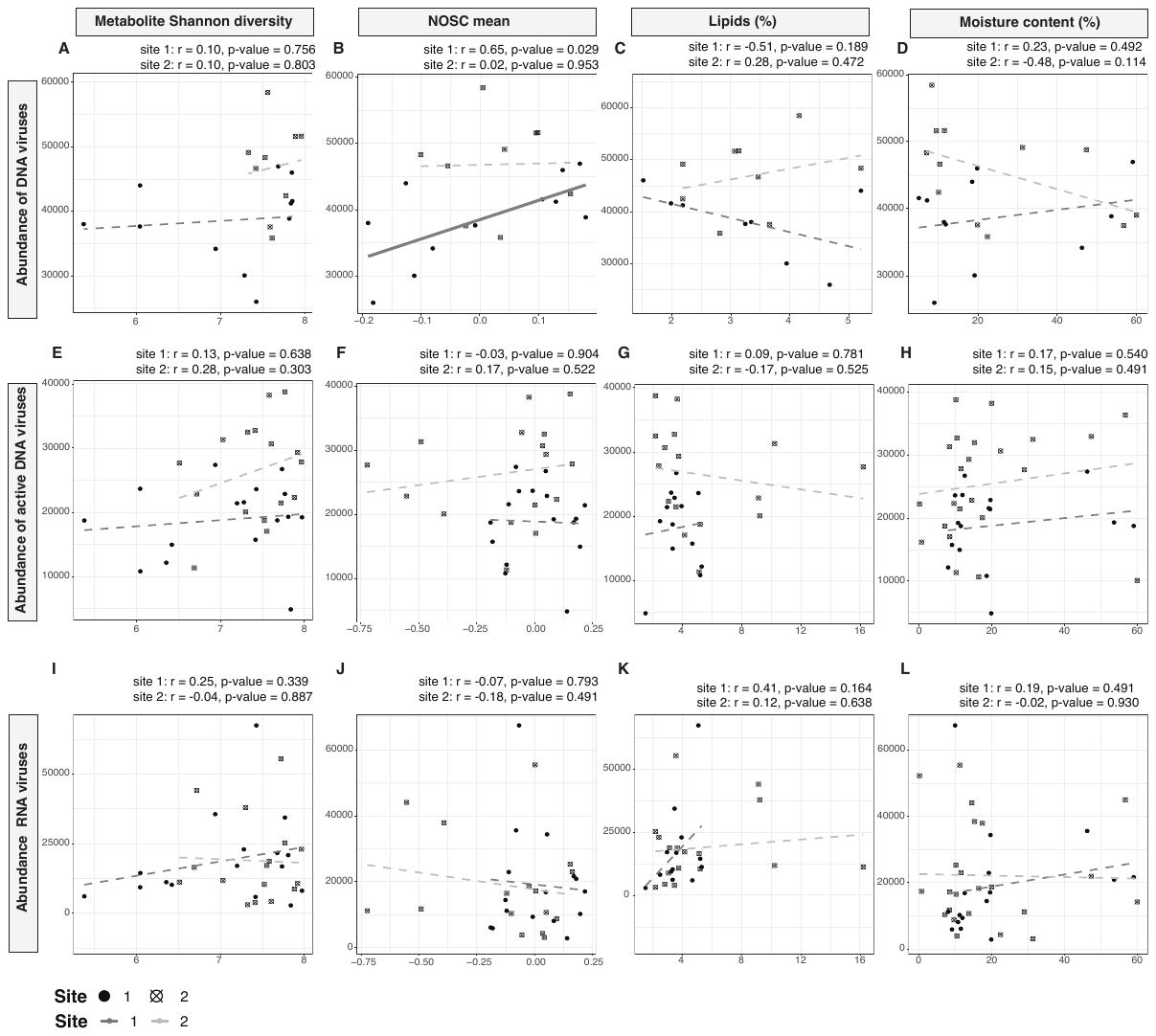

**Supplementary Tables**

**Supplementary Table 1. Drought history of the tropical rainforest in Biosphere 2.**

| **Date (start date)** | **End date** | **Duration (days)** |
| --- | --- | --- |
| 1/16/2000 | 02/11/2000 | 25 |
| 2/18/2000 | 03/16/2000 | 26 |
| 4/13/2002 | 5/13/2002 | 29 |
| 9/23/2002 | 10/28/2002 | 34 |
| 4/21/2003 | 5/6/2003 | 14 |
| 7/23/2014 | 9/29/2014 | 67 |
| 11/1/2015 | 12/2015 | 60 |
| 10/7/2019 | 12/12/19 | 65 |

**Supplementary Table 2. Plant species present in each site.**

| Site 1 | • Ceiba pentandra L.  • Hibiscus tiliaceus L.  • Pachira aquatica Aubl.  • Hibiscus rosa-sinensis  • Elaeis guinensis  • Epipremnum aureum  • Syngonium podophyllum  • Calathea sp.  • Dieffenbachia spp.  • Costus spp.  • Aglaonema crispum |
| --- | --- |
| Site 2 | • Clitoria fairchildiana R.A. Howard  • Ceiba pentandra L.  • Hura crepitans L.  • Pachira aquatica Aubl.  • Hibiscus tiliaceus L,  • Hibiscus rosa sinensis  • Couroupita subsessili  • Musa spp.  • Costus spp.  • Syngonium podophyllum  • Epipremnum aureum  • Heliconia spp.  • Calathea spp |

**Supplementary Table 3. Sporulation factors and resuscitation promoting factors analyzed and their KO numbers (KEGG identification number).**

| **KO** | **Gene name** |
| --- | --- |
| K02490 | spo0F |
| K04769 | spoVT |
| K06283 | spoIIID |
| K06375 | spo0B |
| K06376 | spo0E |
| K06377 | spo0M |
| K06378 | spoIIAA |
| K06379 | spoIIAB |
| K06380 | spoIIB |
| K06381 | spoIID |
| K06382 | spoIIE |
| K06383 | spoIIGA |
| K06384 | spoIIM |
| K06385 | spoIIP |
| K06386 | spoIIQ |
| K06387 | spoIIR |
| K06388 | spoIISA |
| K06389 | spoIISB |
| K06390 | spoIIIAA |
| K06391 | spoIIIAB |
| K06392 | spoIIIAC |
| K06393 | spoIIIAD |
| K06394 | spoIIIAE |
| K06395 | spoIIIAF |
| K06396 | spoIIIAG |
| K06397 | spoIIIAH |
| K06398 | spoIVA |
| K06399 | spoIVB |
| K06401 | spoIVFA |
| K06402 | spoIVFB |
| K06403 | spoVAA |
| K06404 | spoVAB |
| K06405 | spoVAC |
| K06406 | spoVAD |
| K06407 | spoVAE |
| K06408 | spoVAF |
| K06409 | spoVB |
| K06412 | spoVG |
| K06413 | spoVK |
| K06414 | spoVM |
| K06415 | spoVR |
| K06416 | spoVS |
| K06417 | spoVID |
| K07699 | spo0A |
| K08384 | spoVD |

**Supplementary Table 4. Statistical results summarizing how viral community relative abundance in TPM vary based on experimental phase and site.**

| **Response variable** | **Explanatory variable** | **F** | **P** |  | **R^2^** |
| --- | --- | --- | --- | --- | --- |
| DNA virus relative abundance | Phase | 0.49 | 0.62 |  | 0.42 |
|  | Site | 8.79 | 0.008 | Adjusted | 0.26 |
|  | Phase:Site | 1.69 | 0.213 |  | 0 |
| DNA virus community composition | Phase | 1.34 | 0.074 |  | 0.1 |
|  | Site | 3.66 | 0.001 |  | 0.14 |
|  | Phase:Site | 1.07 | 0.349 |  | 0.08 |
| Active DNA virus relative abundance | Phase | 0.45 | 0.645 |  | 0.44 |
|  | Site | 10.12 | 0.006 | Adjusted | 0.27 |
|  | Phase:Site | 0.81 | 0.462 |  | 0 |
| Active DNA virus community composition | Phase | 1.81 | 0.017 |  | 0.11 |
|  | Site | 3.52 | 0.003 |  | 0.07 |
|  | Phase:Site | 0.96 | 0.507 |  | 0.06 |
| RNA virus relative abundance | Phase | 3.92 | 0.015 |  | 0.25 |
|  | Site | 0.12 | 0.731 | Adjusted | 0.11 |
|  | Phase:Site | 0.28 | 0.84 |  | 0 |
| RNA virus community composition | Phase | 3.54 | 0.001 |  | 0.2 |
|  | Site | 1.02 | 0.367 |  | 0.02 |
|  | Phase:Site | 0.73 | 0.657 |  | 0.04 |

**Supplementary Table 5. Results of the calculation of Pearson’s and Spearman correlation between viral relative abundance (RPKM) and soil metabolite properties and moisture content.** Significant results marked in bold font.

| **Variable** | **Variable** | **method** | **R/rho** | **P** |
| --- | --- | --- | --- | --- |
| DNA virus relative abundance | Metabolite Shannon diversity | Pearson | 0.49 | **0.02788** |
|  |  | Spearman | 0.51 | **0.02272** |
|  | NOSC mean | Pearson | 0.46 | **0.04116** |
|  |  | Spearman | 0.35 | 0.1303 |
|  | Lipids (%) | Pearson | 0.20 | 0.2195 |
|  |  | Spearman | -0.016573 | 0.9447 |
|  | Moisture content (%) | Pearson | -0.111219 | 0.6134 |
|  |  | Spearman | -0.164032 | 0.4528 |
| Transcriptionally active DNA virus relative abundance | Metabolite Shannon diversity | Pearson | 0.3372 | 0.1856 |
|  |  | Spearman | 0.308824 | 0.2273 |
|  | NOSC mean | Pearson | 0.325413 | 0.2025 |
|  |  | Spearman | 0.428922 | 0.08733 |
|  | Lipids (%) | Pearson | 0.221654 | 0.3926 |
|  |  | Spearman | -0.374233 | 0.1389 |
|  | Moisture content (%) | Pearson | 0.170601 | 0.4721 |
|  |  | Spearman | 0.377444 | 0.1017 |
| RNA virus relative abundance | Metabolite Shannon diversity | Pearson | 0.184851 | 0.4628 |
|  |  | Spearman | 0.188855 | 0.4514 |
|  | NOSC mean | Pearson | 0.248635 | 0.3198 |
|  |  | Spearman | 0.23839 | 0.3393 |
|  | Lipids (%) | Pearson | -0.556039 | **0.01657** |
|  |  | Spearman | -0.243802 | 0.3296 |
|  | Moisture content (%) | Pearson | 0.707587 | **0.000333** |
|  |  | Spearman | 0.555844 | **0.009972** |
